# Systematic benchmark of demultiplexing workflows for direct RNA sequencing

**DOI:** 10.64898/2026.10.03.756430

**Authors:** James William Bryson, Lena Bjørn Johansson, Anila George, Junyu Chen, Jamie Yam Auxillos, Sarah Rennie

## Abstract

Direct RNA nanopore sequencing offers the capacity to directly profile native RNAs, enabling detection of diverse epitranscriptomic modifications based on current changes as the RNA passes through a pore. Innovations in chemistry, hardware and bioinformatic tools have increased sequencing capacity and accuracy, creating demand for sequencing multiple samples within individual runs. Two recent solutions enable barcoding and demultiplexing of samples, but independent comparisons of their performance is limited. Here we benchmark WarpDemuX and SeqTagger using orthogonal ground truths derived from species identity and matched IVT spike-ins across four RNA004 flow cells. Both achieve high demultiplexing precision but differ in their trade-offs in precision and retention. SeqTagger provided strong performance at its recommended settings, whilst WarpDemuX performance varied by model, barcode and read quality. IVT-derived score cutoffs allowed dataset-specific calibration and increased WarpDeMux assignment yield, although precision achieved on IVTs was not consistently reproduced in species reads. These results support matched IVTs as internal controls for guiding choice of demultiplexing settings, while demonstrating the need to validate demultiplexing precision in biological RNA populations.

## Introduction

Oxford Nanopore direct RNA sequencing (DRS) is a rapidly expanding transcriptomic technology due to its capability to directly sequence full-length native RNA molecules by measuring changes in ionic current as RNA molecules pass through a nanopore (Garalde et al., 2018). Unlike cDNA-based sequencing methods, DRS does not require PCR amplification, allowing for preservation of molecular features such as poly(A)-tail information and signal changes associated with epitranscriptomic modifications (Garalde et al., 2018; Workman et al., 2019). Successive improvements in sequencing chemistry and basecalling, including introduction of RNA004, have increased read accuracy and depth (Liu-Wei et al., 2024). This in turn makes DRS attractive for large-scale studies of post-transcriptional regulation. In such studies, multiplexing of samples on individual flow cells can reduce sequencing costs and limit confounding due to batch and run effects. However, DRS multiplexing has historically been technically demanding because barcode information encoded in the DNA reverse-transcription adaptor must be recovered from signal generated under direct RNA sequencing conditions.

Early solutions such as DeePlexiCon established proof of principle for demultiplexing of DRS libraries (Smith et al., 2020). DeePlexiCon incorporated custom DNA barcodes into the reverse-transcription adaptor, and transformed the raw nanopore signal into images subsequently classified using convolutional neural networks (Smith et al., 2020). More recent approaches have extended this principle, with WarpDemuX and SeqTagger demonstrating high-accuracy demultiplexing across larger barcode sets (Pryszcz et al., 2025; van der Toorn et al., 2025). WarpDemuX works independently of basecalling (van der Toorn et al., 2025). It first finds the adaptor region in the raw signal and summarises the barcode-associated signal as a short fingerprint, then compares these fingerprints using dynamic time warping and classifies the reads with a support vector machine. WarpDemuX also considers barcode design in signal space, by choosing barcodes based on how separable their predicted nanopore signals are, rather than only how different their DNA sequences are. Its models for RNA004 include WDX4 supporting four predefined barcodes, and the ten barcode model WDX10.

SeqTagger takes a different route. It first extracts the part of the signal that contains the barcode, then directly basecalls the barcode sequence using a custom Bonito-trained model. The decoded barcode is aligned to a known barcode set with minimap2 (Li, 2018), and low-confidence assignments can be filtered using barcode base-quality scores. In this way, SeqTagger shifts the task from raw-signal classification to barcode-specific basecalling and mapping, supporting models with up to 96 barcodes (Diensthuber et al., 2026; Pryszcz et al., 2024). However, independent benchmarks comparing these approaches under conditions involving heterogeneous samples remain limited.

As DRS is used to measure an increasing range of RNA features, often requiring well-calibrated deep learning tools, suitable reference controls are needed to assess the reliability of these measurements. In vitro transcribed (IVT) RNA spike-ins provide controls for measurement bias, transcript structure and isoform quantification in RNA sequencing (Hardwick et al., 2016; Jiang et al., 2011). In DRS, synthetic controls have supported poly(A)-tail estimation using transcripts with defined tail lengths (Chang et al., 2025), and modification detection using unmodified IVT RNA as a comparator (Leger et al., 2021; Liu-Wei et al., 2024). These applications illustrate how RNA features can be anchored to independent references within individual study designs (The International Human RNome Project Consortium, 2025). IVT spike-ins with distinct known sequence identities could similarly be used to assess and calibrate demultiplexing with individual sequencing runs, although the full utility of this approach remains inadequately explored.

Here, we systematically benchmark DRS multiplexing performance across synthetic and biologically distinct RNA mixtures. We used IVT spike-ins with defined sequences together with transcriptomes from four different species to establish barcode-independent reference identities across two DRS runs per workflow (WarpDemuX and SeqTagger). We assessed precision, in-panel read retention and off-panel assignment under the published workflow settings, showing that both approaches generally achieve high in-panel precision, with differences in their performance strongly associated with read qualities and selected barcode confidence thresholds. We used IVTs to calibrate thresholds and tested their transfer to species reads within and across runs, alongside their effect on assignment yield across all reads. Together, these analyses provide an independent benchmark of demultiplexing approaches in heterogeneous biological samples and evaluate use of matched IVT controls as a practical basis for selecting demultiplexing settings.

## Results

### Study design establishes ground-truth read identities independently of barcode prediction

We designed a benchmark for direct RNA sequencing demultiplexing methods using ground-truth read identities that could be established independently of demultiplexing. Poly(A)+ RNA derived from each of four different species (*A. thaliana*, mouse, human and *S. cerevisiae*) was paired with one of four enzymatically tailed IVT spike-ins with distinct sequence identities and four barcoded samples were loaded per flow cell (**FC1 - FC4, Figure 1A,B**). WarpDemuX libraries were sequenced on FC1 and FC4 and SeqTagger libraries were sequenced on FC2 and FC3. Barcode assignments to the fixed species - IVT pairs were rotated between the two runs for each workflow to provide observations of the same RNA origin under different barcode assignments (**Figure 1B**). In this way, the two WarpDemuX models WDX4 and WDX10 could be evaluated on the same read sets with two sources of reference identity, whilst SeqTagger could be assessed similarly on separate flow cells.

**Figure 1.**
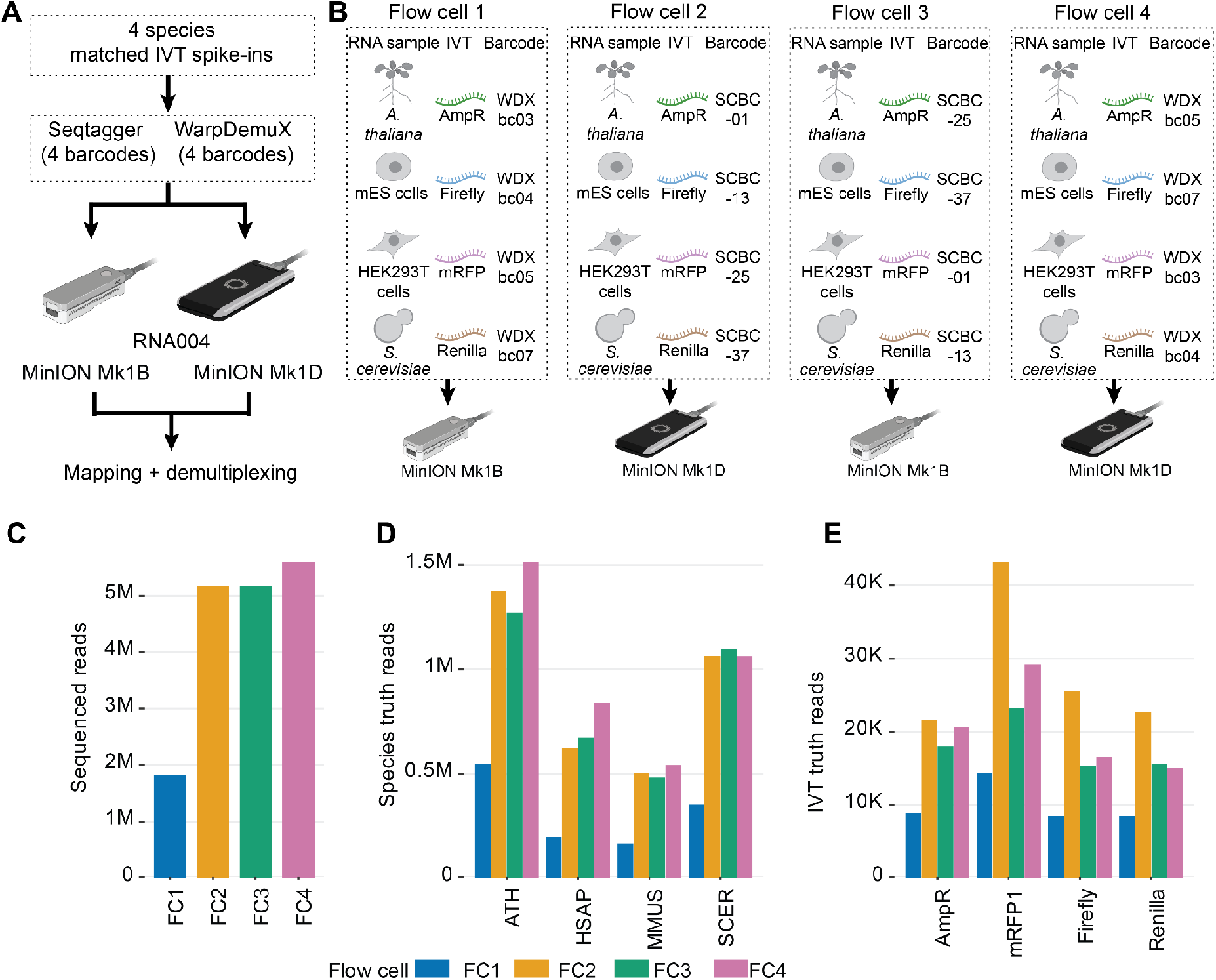
Experimental design and independent mapping reference sets. **(A)** Library design with four species and four distinct IVT RNAs. **(B) S**pecies - IVT pairs and barcode assignments in each flow cell. **(C)** Total number of basecalled reads per flow cell. **(D)** Number of reads distinctly mapping to a single species with MAPQ >= 60 and at least 80% of the read aligned, per flow cell. **(E)** As in (D) but for IVTs. WDX4 and WDX10 use FC1 and FC4, SeqTagger uses FC2 and FC3. The reference reads in (D) and (E) define the ground-truth sets used in subsequent analyses. ATH, *Arabidopsis thaliana*; HSAP, *Homo sapiens*; MMUS, *Mus musculus*; SCER, *Saccharomyces cerevisiae*.

Sequencing yielded a total of 17,747,356 reads, ranging from 1.8 million reads in FC1 to 5.6 million reads in FC4 (**Figure 1C**), which were subsequently basecalled using Oxford Nanopore’s Dorado software (Oxford Nanopore Technologies, 2026a). Mean read-quality and read-length distributions were consistent across all flow cells (**Supplementary Figure 1A,B**). We established ground-truth read sets by aligning reads to a combined reference comprising the four species genomes and the four IVT sequences, requiring a mapping quality (MAPQ) of at least 60 and at least 80% of each read to align. We also examined the sensitivity of the reference read counts to the selected MAPQ threshold, which were largely invariant to modest deviations (**Supplementary Figure 1C**). The alignment requirement excluded 3.6% of the species reads and 3.8% of the IVT reads passing the MAPQ score cut off (**Supplementary Figure 1D**). The criteria retained a total of 12,317,897 species reads and 307,023 IVT reads, with substantial differences in the abundance of each origin observed within the flow cells (**Figure 1D, E**). Subsequent benchmarking analyses used these sets as fixed references with known origin.

### Default demultiplexing workflows achieve high precision, with lower retention for specific reads than IVT controls

We evaluated three models for demultiplexing: WarpDemuX *WDX4*, which was trained on the same four-barcode set used in this study, WarpDemuX *WDX10*, which was trained on a set of ten barcodes including these four, and SeqTagger *b96_RNA004*, which distinguishes amongst 96 barcodes, of which four were used here. WDX10 and SeqTagger therefore have six and 92 off-panel classes, respectively, corresponding to barcodes absent from the libraries. WDX4 has no off-panel classes because all four of its modelled barcodes were loaded. SeqTagger assignments were filtered at the recommended barcode base quality-score (baseQ), requiring baseQ >= 50. WarpDemuX was evaluated using the calls passing its nominal 99% settings, which apply per-barcode cutoffs to its internal confidence score, defined as the difference between the best score and second-highest class scores.

For each model, final assignments were classified as in-panel, off-panel or not assigned (**Figure 2A**), with confusion matrices showing within-panel misassignments (**Supplementary Figure 2**). The proportion of reads assigned in-panel to a loaded barcode for IVTs was overall high, although there were noticeably higher rates for Seqtagger (95.5 - 95.7%) and WDX4 (95.7 - 95.8%) than for WDX10 (87.4 - 88.2%). Within each method, the in-panel proportion was systematically lower for species compared to IVT **(Figure 2A)**. There were very few off-panel calls in either IVT or species, and the reduced in-panel read assignment rates observed in species mainly reflected an increase in unassignment rates (8.0 - 13.0% in SeqTagger, 8.4 - 9.9% in WDX4 and 16.8 - 19.0% in WDX10) (**Figure 2A, Supplementary Figure 2**).

**Figure 2.**
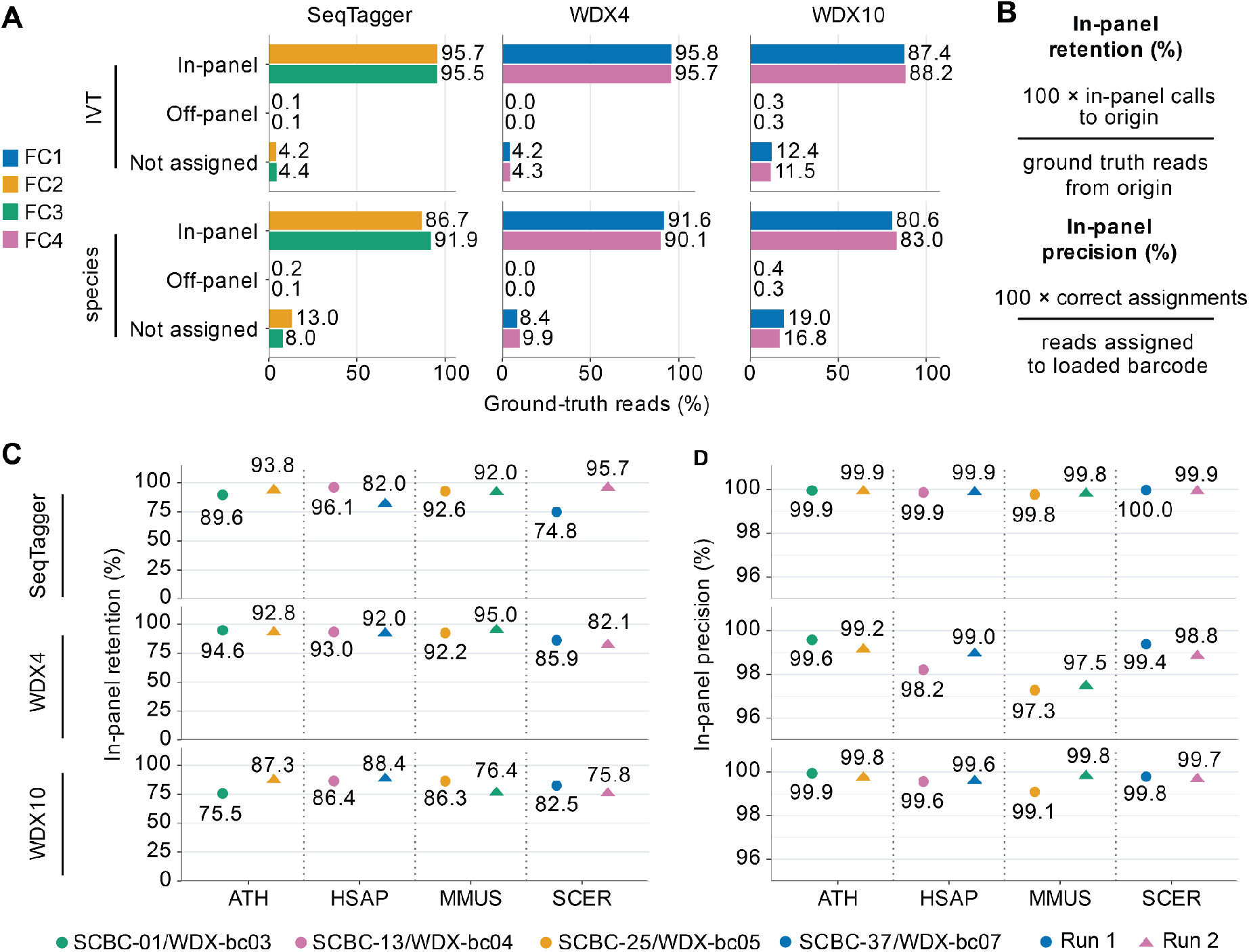
Barcode assignments under the intended workflows. **(A)** In-panel, off-panel and not-assigned read fractions within each species or IVT reference set and flow cell. **(B)** Definitions of in-panel retention and precision. **(C)** In-panel retention for species reference reads according to read origin, including incorrect assignments to loaded barcodes. **(D)** In-panel precision for each assigned barcode, labelled by its corresponding species. Colours indicate barcode identities, circles and triangles denote run 1 and run 2, respectively. SeqTagger calls are based on baseQ>=50 and WDX calls are shown at nominal 99% settings, defined in the original publication.

For a given origin, in-panel retention was calculated as the proportion of its ground-truth reads assigned to any loaded barcode, and in-panel precision was calculated for each barcode as the fraction of assigned reads that came from the expected origin **(Figure 2B)**. In-panel retention varied across species and barcode assignments **(Figure 2C)**. For example, SCBC-37 resulted in the two lowest values for SeqTagger (82.0% and 74.8%), whilst WDX10 retention for bc03 was approximately 76% in both runs, despite in both cases the barcode being assigned to different species. All three models achieved high in-panel precision, with SeqTagger consistently close to 100% and greater barcode to barcode variation observed for WDX4 (**Figure 2D**). We additionally noted that reads mapping to mouse generally had lower in-panel precision compared to other species, suggesting that relative abundance might explain some of the observed differences in in-panel precision.

### Read quality is associated with variation in demultiplexing performance

We next examined whether lower retention in species was a uniform feature or associated with specific read subsets. In-panel retention was strongly associated with mean RNA read quality, a measure which was based on the molecule itself and distinct from the barcode baseQ used to filter SeqTagger calls (**Figure 3A**). Overall, IVT and species derived reads had very similar quality score distributions across a set of defined bins (**Supplementary Figure 3A**). Notably, amongst species reads with mean RNA quality scores below 10, retention was as low as 53% for SeqTagger, 72% for WDX4 and 52.3% for WDX10, and increased strongly for scores up to around 20. SeqTagger maintained a high precision even in the lowest quality bin (**Figure 3B**), whilst precision was reduced in WDX4 and WDX10 below a quality score of 15.

**Figure 3.**
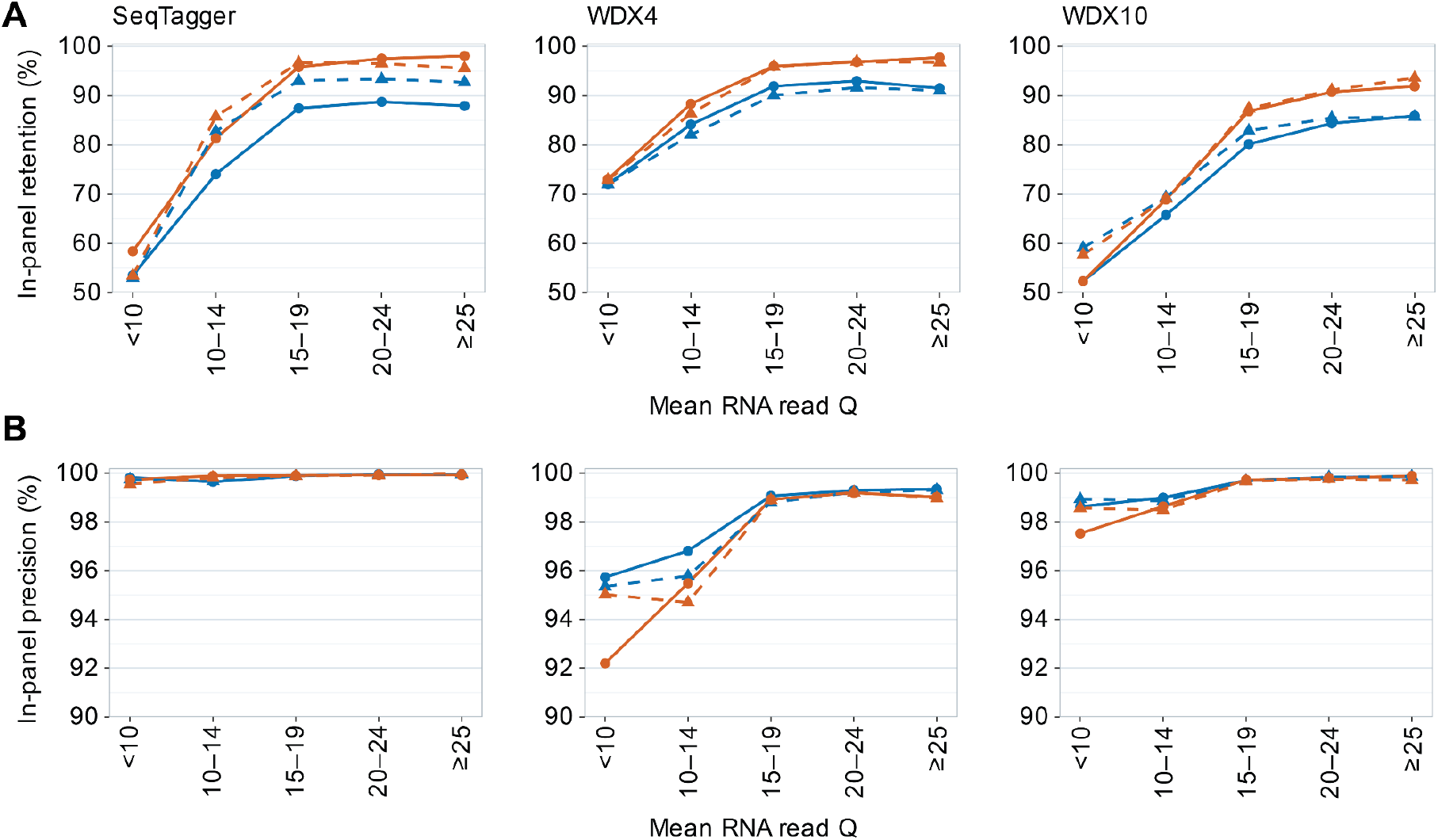
Association between RNA read quality and demultiplexing performance. **(A)** In-panel retention for reads binned according to mean RNA read quality, for species (blue) and IVT (orange) reference reads. Solid lines with round circles denote run 1 and dashed lines with triangles denote run 2. **(B)** As in (A) but using in-panel precision. Metrics are based on published workflow settings and all ground-truth reference reads within each quality bin.

In addition to read quality we investigated whether the ability to call poly(A) tails from the reads by Dorado was associated with demultiplexing outcomes. Although all RNAs should be poly(A)+, we observed large barcode to barcode differences in the percentage of reads with a detectable tail, which was independent of species and IVT origin (**Supplementary Figure 3B**). For species, modestly lower retention rates were observed in all methods for reads without a recorded poly(A) tail, a pattern which was not observed for IVT reads, and precision was only modestly affected for WarpDemuX (**Supplementary Figure 3C,D**). Notably, neither quality score nor tail estimation was able to explain the higher retention rates of IVT reads compared to species. Thus, heterogeneous read properties were associated with demultiplexing performance but did not account for all variability.

### Varying barcode score thresholds reveals method-specific precision-recall trade-offs

Demultiplexing methods aim to maximise read retention whilst maintaining high precision in order to minimise cross-sample misassignment. SeqTagger relies on a single, common baseQ cutoff across barcodes, whereas WarpDemuX uses per-barcode confidence thresholds calibrated for a target precision. To examine the effect of these thresholds, we generated precision - retention curves for each barcode by varying its score cutoff **(Figure 4A)**. Because precision depends on the relative abundance of each species within a flow cell, we weighted reference reads so that each species contributed equally within its flow cell, allowing for the barcodes to be compared without the results being dominated by variation in species abundance (corresponding curves based on unweighted reads are shown in **Supplementary Figure 4A**).

**Figure 4.**
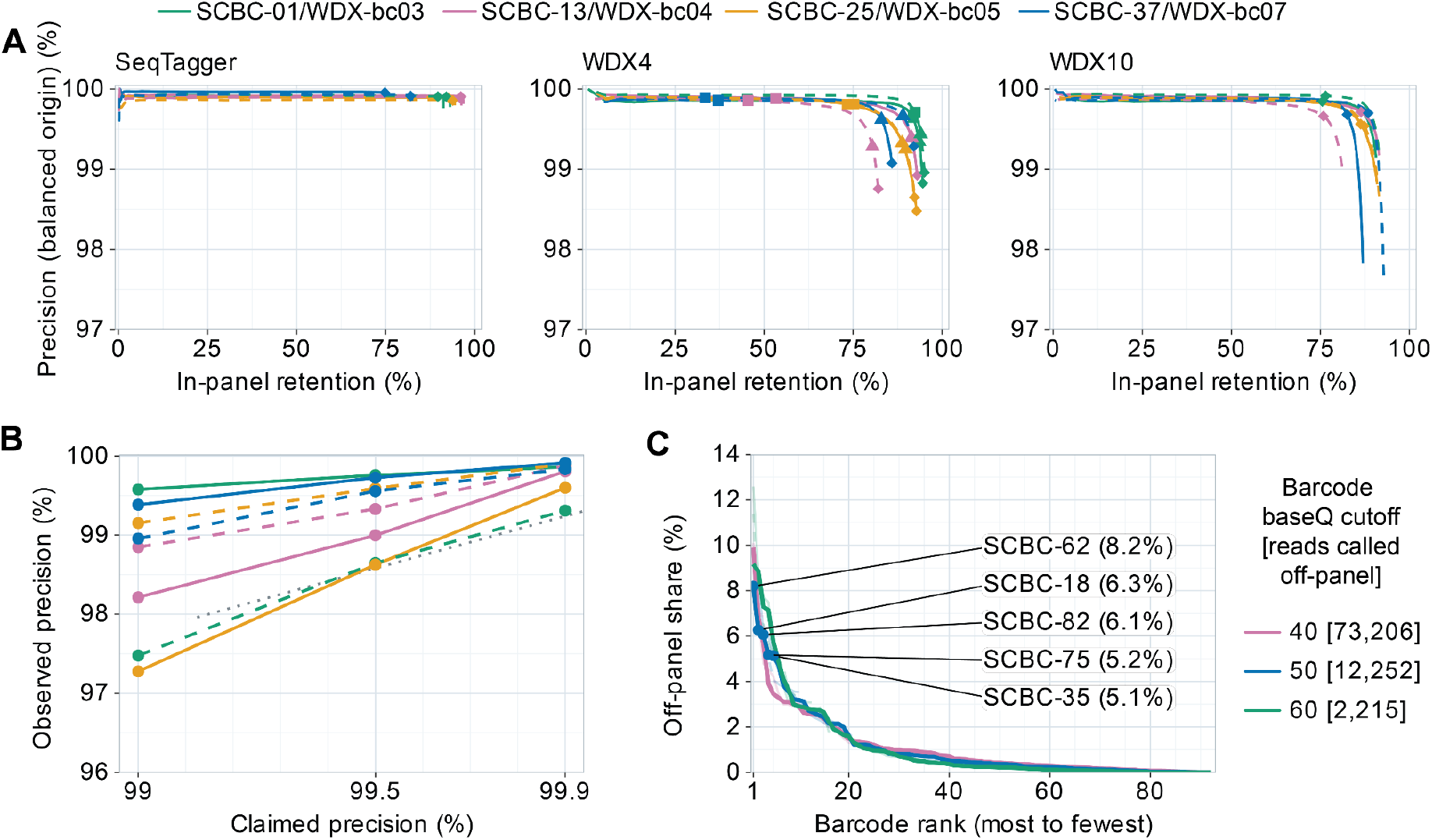
Barcode-specific precision–retention curves, nominal targets and off-panel assignments. **(A)** Species precision versus in-panel retention, after weighting each origin equally within its flow cell. All reference reads are retained; each species contributes one-quarter of the total weight. Each curve varies the score cutoff for one barcode (baseQ for SeqTagger, confidence score for WarpDemuX), with other barcodes kept at their default settings. Colours indicate different barcodes and solid and dashed lines denote runs 1 and 2. Symbols mark the default settings and the stricter WDX4 targets. **(B)** Observed WDX4 species precision versus the nominal target, separated by barcode and run. Metrics are based on unweighted read counts at the supplied 99%, 99.5% and 99.9% settings. The dashed identity line indicates equal observed and nominal precision. **(C)** Ranked shares of SeqTagger off-panel species calls at baseQ cutoffs of 40, 50 and 60, with species and runs weighted equally. Thick curves combine the two runs and faint curves show individual runs. Bracketed legend values give the total observed off-panel read counts. Labels identify the five most frequent off-panel barcodes at baseQ >=50.

SeqTagger maintained approximately 99.9% in-panel precision across its baseQ cutoff range **(Figure 4A)**. For both WarpDemuX models, precision decreased as the cutoffs became less stringent, with the largest changes near the highest attainable retention. The extent of this decline varied between barcodes, showing that the additional read retention obtained by relaxing the cutoff came at different costs to precision.

We next examined how closely the precision achieved using the original WDX4 thresholds matched their nominal targets. Applying the original WDX4 per-barcode thresholds for targets of 99%, 99.5% and 99.9%, we observed increasing precision at more stringent settings for both IVT and species reads. However, the precision achieved at each target varied by barcode and run and often fell below the nominal value **(Supplementary Figure 4B, Figure 4B)**.

Finally, we analysed the distribution of off-panel calls. For WDX10, the numbers of calls to individual unused barcodes varied between runs (**Supplementary Figure 4C)**. Finally, we analysed the distribution of SeqTagger off-panel calls, which showed they were concentrated in a subset of unused barcodes **(Figure 4C)**. Furthermore, increasing the baseQ cutoff from 40 to 50 reduced the number of off-panel species calls across the two runs from 73,206 to 12,252 and a cutoff of 60 reduced this further to 2,215. Thus, although SeqTagger’s in-panel precision remained largely unchanged across cutoffs, stricter baseQ filtering substantially reduced assignments to barcodes absent from the library.

### IVT controls support dataset-specific calibration of demultiplexing thresholds

Next, we asked whether the relationship between barcode confidence and precision could be estimated from the spiked-in IVT controls, providing an internal, dataset-specific calibration. For each flow cell, we selected four barcode-specific WarpDemuX cutoffs or one SeqTagger baseQ cutoff using IVT ground-truth reads alone, to meet the specified IVT targeted precision (**Figure 5A)**. The selected cutoffs (**Supplementary Table 2)** were then applied unchanged to species reads within the same flow cell (cis) or in the other run (trans).

**Figure 5:**
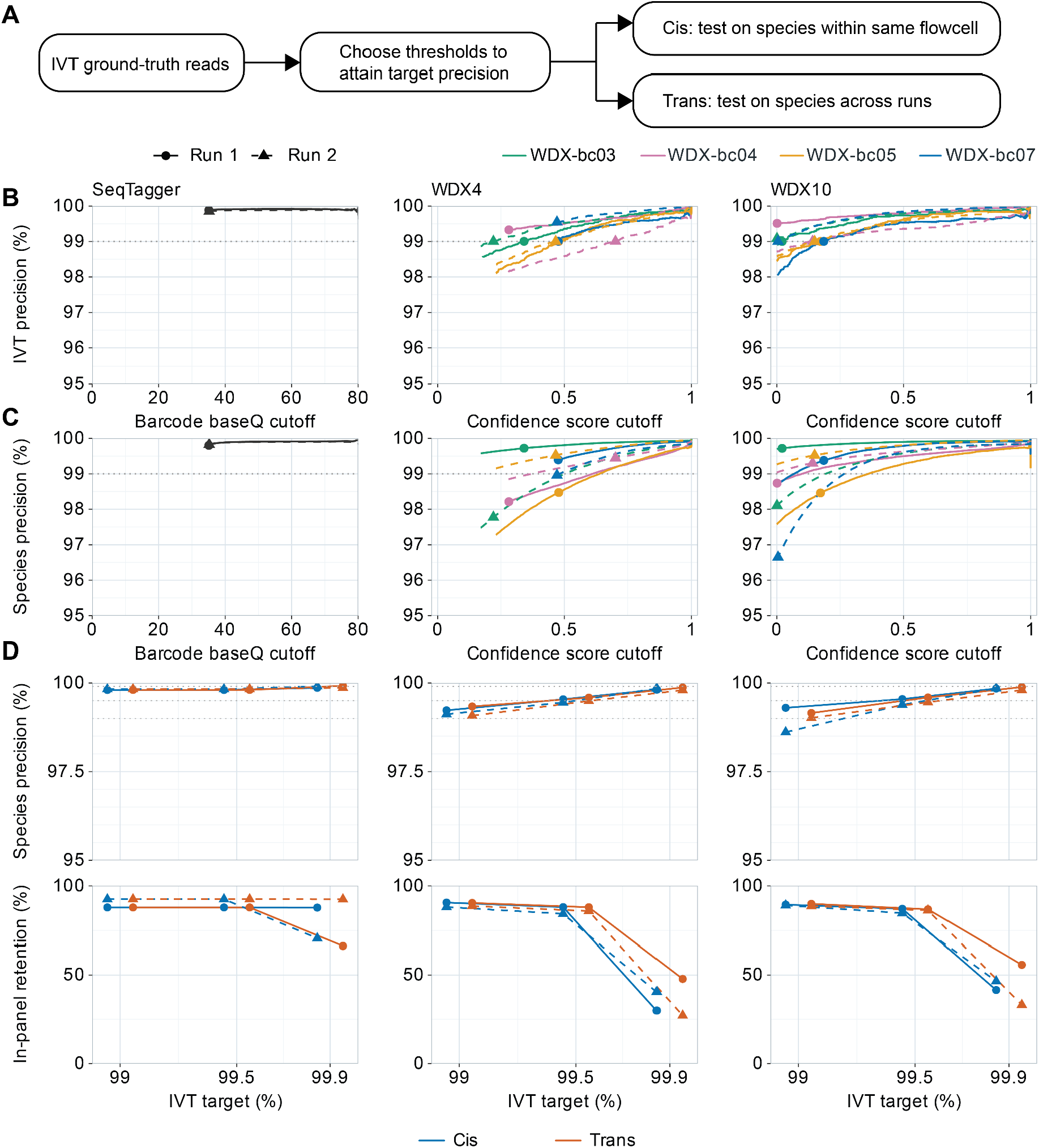
IVT-based calibration and species evaluation within and across runs. **(A)** Schematic illustrating selection of score cutoffs using IVT reads, followed by evaluation on species reads within the same flow cell (cis) or in the other run (trans). **(B)** IVT precision across score cutoffs, with symbols marking the selected 99% target settings. SeqTagger uses a single cutoff; each WarpDemuX barcode is calibrated separately. **(C)** Species precision across score cutoffs, with symbols marking the IVT-selected cutoffs from B applied within the same flow cell. Colours identify WarpDemuX barcodes; solid and dashed lines denote runs 1 and 2. **(D)** Species precision (top) and in-panel retention (bottom) at IVT targets of 99%, 99.5% and 99.9%, for cis (blue) and trans (orange) calibration. Circles and triangles denote runs 1 and 2, respectively.

For both 99% and 99.5% precision targets, SeqTagger had an in-panel IVT precision of 99.89% in FC2 and 99.85% in FC3 without applying a baseQ filter, where the lowest observed scores of 35.1 and 35.2 amongst in-panel IVT assignments from FC2 and FC3 respectively, was selected (**Figure 5B, Supplementary Figure 5A)**. This resulted in a species precision of 99.80 - 99.83% (**Figure 5C, Supplementary Figure 5B)**. For WarpDemuX, selected scores per barcode varied and resulted in a within-flow cell species precision close to the expected 99% (99.12–99.23% for WDX4 and 98.61–99.30% for WDX10) (**Figure 5D)**. For all methods, including the IVT target to 99.5% improved species precision came at a cost to retention (**Figure 5D**). Interestingly, across-run transfer (trans) produced similar aggregate performance (**Figure 5D**).

Finally, IVT subsampling yielded successful calibrations in all 100 draws at each tested size from 2,000 reads upwards. Larger samples generally reduced variation in the resulting species precision and retention (**Supplementary Figure 5C**), showing that calibration became more stable as the number of IVT reads increased.

### IVT calibration increases WDX10 assignment yield and overlap with WDX4

We next applied the default and the 99% IVT-calibrated settings to all reads, including those outside of the mapping ground truth reference sets (**Figure 6A**). WDX10 assigned approximately 5.6 million reads to loaded barcodes at default settings, increasing to 6.1 million under cis calibration and only 20,000 fewer under trans calibration, corresponding to a gain of 8.83 - 9.18%. SeqTagger assignments increased by approximately 1.66% across FC2 and FC3, whilst WDX4 assignments decreased by 1-2% (**Figure 6A**).

**Figure 6.**
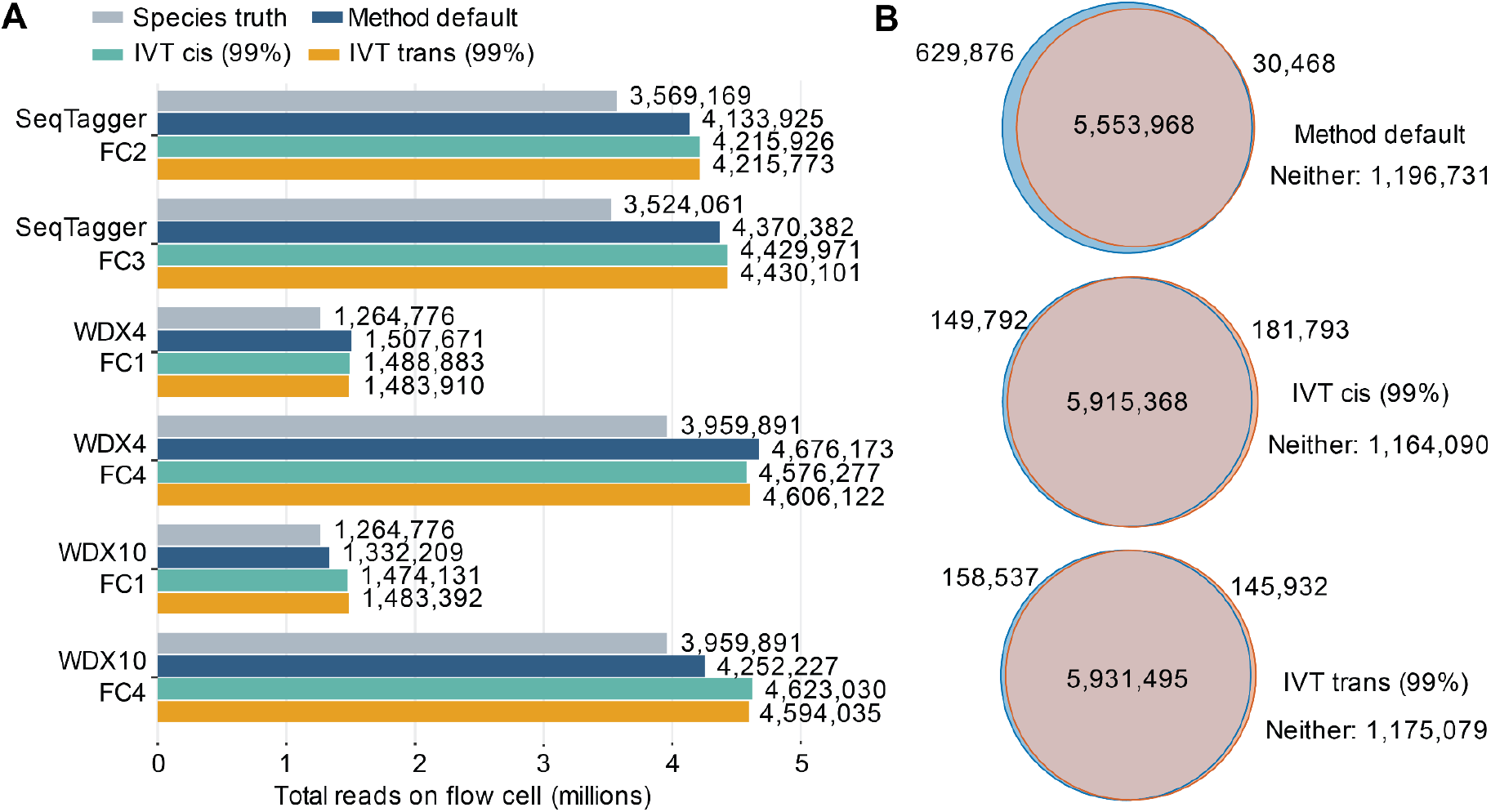
Assignment yield across all reads. **(A)** Reads assigned to loaded barcodes at workflow defaults, or at the 99% target IVT-calibrated cis or trans settings, where SeqTagger uses a baseQ score of zero under the calibrated setting. The grey bars show the number of species ground-truth reads, whilst the other bars expand to calling across the full read set regardless of established ground-truth. **(B)** Venn diagrams for WDX4 (blue) and WDX10 (orange) assigned-read sets pooled across FC1 and FC4, separately for default, cis and trans settings. The overlap counts reads assigned by both models. Counts of reads that were not assigned by either model are designated “Neither” on the figure.

Finally, pooling the two WarpDemuX runs and comparing the WDX10 and WDX4 assigned-read sets showed that, using default settings, 89.37% of reads assigned by either model were assigned by both (**Figure 6B**). After calibration, the overlap increased to 94.69% under cis calibration and 95.12% under trans calibration. Calibration therefore increased agreement between the models in which reads were retained for analysis.

## Discussion

Our results complement the original studies by independently evaluating demultiplexing jointly across matched synthetic IVT controls and, importantly, heterogeneous RNA samples from four species. These samples capture variation in RNA composition and read quality relevant to the biological samples that users seek to multiplex. Overall, both workflows achieved high precision under their published settings, although performance varied across barcodes, RNA populations and score thresholds. WDX10 was more selective than WDX4 for the same read sets, whilst SeqTagger maintained high precision across read-quality bins. The consistently higher retention of IVTs demonstrates the value of evaluating synthetic controls alongside native RNA, because the species samples revealed differences in the handling of lower-quality reads that are important for predicting how much data will remain available for biological analysis.

Because the barcodes used in a run are known, off-panel calls can be identified and excluded. We therefore focussed our metrics on in-panel retention and precision, reflecting the practical workflow of separating reads by barcode after removing unassigned and off-panel reads. SeqTagger with its recommended baseQ threshold provides a useful balance of precision and retention without additional tuning. For WarpDemuX, per-barcode confidence thresholds offer flexibility in choosing this balance, and the precision–retention curves show the read loss associated with stricter filtering. The variation in observed precision at the nominal WDX4 targets also highlights the value of checking performance within individual datasets.

Matched IVTs provide a practical way to make this assessment and select thresholds using controls sequenced alongside each sample. Their distinct sequences allow barcode assignments to be checked independently, and the selected thresholds can then be applied to native RNA from the same library. WDX10 benefited most from this calibration, with approximately 9% more reads assigned to loaded barcodes and greater overlap with the reads selected by WDX4. The comparison with species references shows how these gains in retention relate to precision on heterogeneous biological RNA.

The cis and trans comparisons provide an initial test of whether calibration can be reused between runs. The runs were performed one week apart, making this a relevant test of short-term transfer within one laboratory. Similar performance between cis and trans calibration at the 99% and 99.5% targets supports reuse of thresholds between closely matched runs. At the 99.9% target, retention was more sensitive to the calibration run, and stringent filtering removed a substantial fraction of reads. This illustrates the importance of choosing a precision target appropriate to the study and assessing the associated loss in sequencing depth.

Poly(A) tail measurements may also be affected by the use of custom DNA barcode sequences. For both SeqTagger and WarpDemuX, Dorado did not estimate tail length for a subset of reads, even though the input population was poly(A)+ and the DRS protocol relies on polyadenylated RNA for attachment of the reverse transcription adaptor. However, we note that our subsequent runs using the same SeqTagger barcodes on the high-throughput PromethION RNA004 flow cells showed consistently high poly(A) tail call rates (data not shown). Our results motivate further testing of how barcode design, read quality and sequencing platform affect poly(A)-tail estimation.

Some limitations should be noted. Four barcodes were loaded throughout, so all three configurations were tested at four-plex and two of them well below capacity. With more barcodes in use, a misassignment to a barcode that is currently unused could be counted as an assignment to another sample, changing the pattern of cross-sample errors. Future designs could distribute the same sample panel across multiple flow cells to reduce confounding between samples and runs and assess the variability we observed in species read counts per flow cell. The ground-truth reference sets were based on read mapping and may contain errors despite the stringent cutoffs. In addition, WarpDemuX was largely evaluated at its default settings, apart from the confidence score thresholds. Further optimisation of other settings may improve performance.

Finally, Oxford Nanopore now offers the Direct RNA Barcoding Kit 24 (SQK-DRB004.24), providing another, officially supported, multiplexing option. A comparison of preparation efficiency, usable yield, accuracy and cost with the community-based methods evaluated here would help users select between approaches. In addition, the current protocol of this kit discourages mixing native poly(A)+ RNA and poly(A) tailed IVTs (Oxford Nanopore Technologies, 2026b), so its evaluation may require an adapted spike-in design.

## Conclusion

In conclusion, matched IVT spike-ins and species ground-truth references provide complementary measures for benchmarking DRS demultiplexing performance. We have shown that current methods perform well overall, with trade-offs between assignment yield and precision. Matched IVTs offer useful internal controls for selecting demultiplexing thresholds, assessing their consistency between runs and improving read retention. Evaluating these controls alongside heterogeneous biological samples makes the benchmark directly relevant to multiplexed transcriptomic studies.

## Materials and Methods

### Samples, spike-ins and pool design

Libraries were prepared from four species: Arabidopsis thaliana, Homo sapiens, Mus musculus and Saccharomyces cerevisiae (RNA extraction and poly(A) selection described below), alongside four unmodified in vitro transcribed (IVT) spike-in transcripts: mRFP1, AmpR, Firefly luciferase and Renilla luciferase (template construction, IVT synthesis and polyadenylation steps described below). Because each IVT is synthesised and barcoded as a separate reaction, every spike-in read’s barcode is known by design.

During library preparation, each sample and corresponding IVT was barcoded with one barcode from each of two sets. WarpDemuX barcodes used bc03, bc04, bc05 and bc07, and SeqTagger barcodes used SCBC-01, SCBC-13, SCBC-25 and SCBC-37. The libraries were prepared with each of the four species representing 24.5% of the library and each of the four IVT spike-ins representing 0.5% by design (described further in the Nanopore DRS library preparation and sequencing section). Between the two replicates the barcode assigned to each species–IVT pair was changed, so that each pair was observed under two barcodes.

### In-vitro transcriptions

Four IVT sequences of comparable length (0.8-1kb), derived from Firefly luciferase, Renilla luciferase, AmpR, and mRFP1, were selected as RNA standards. Their sequences were screened by BLAST to confirm no significant sequence similarity to the transcriptomes of the four selected species (Altschul et al., 1990). Templates were generated by PCR amplification using primers incorporating a T7 promoter upstream of each sequence. *In vitro* transcription was performed using the Promega RiboMAX Large Scale RNA Production System (Promega P1300) according to the manufacturer’s protocol. Following transcription, template DNA was removed by addition of 1 μL RQ1 DNase and incubation for 15 min then purified using the TRIzol Plus RNA Purification Kit (Thermo 12183555) according to the manufacturer’s protocol. RNA products were quantified using the Nanodrop One (Thermo Fisher) and assessed using an Agilent RNA Pico Bioanalyze. The IVT transcripts were then polyadenylated in 25 μL reactions containing 5× poly(A) polymerase buffer, 25 mM ATP, poly(A) polymerase (Thermo 74225Z25KU), RNase inhibitor, and incubated at 37C for 15 minutes then purified using the Zymo RNA Clean and Concentrator (Zymo R1014), quantified using the Nanodrop One (Thermo Fisher) and assessed for polyadenylation using an Agilent RNA Pico Bioanalyzer.

### Transcriptome purifications

#### Homo sapiens RNA

HEK293T cells were cultured in DMEM (Thermo 11995065) supplemented with 10% fetal bovine serum (Thermo A5256801) and 1% Penicillin/Streptomycin (Thermo 15140122). Cells were grown at 37°C in a humidified incubator with 5% CO_2_ and passaged twice a week once confluency reached 70-80%. RNA was extracted using the TRIzol Plus RNA Purification Kit (Thermo 12183555) according to the manufacturer’s protocol and eluted into DNA LoBind tubes (Eppendorf 0030108.418).

#### Saccharomyces cerevisiae RNA

*Saccharomyces cerevisiae* (BY4741) cultures were grown in YPD media in shaking conditions at 180 rpm at 30°C overnight then, diluted to OD600 0.2, and harvested in mid-log phase at OD600 ~0.6. Cells were pelleted, washed once in PBS, and treated with Zymolyase Ultra (Zymo E1007-2) and Y1 buffer to digest the cell wall. Following incubation at 30°C for 30 minutes, cells were pelleted and RNA was extracted using the TRIzol Plus RNA Purification Kit (Thermo 12183555) according to the manufacturer’s protocol and eluted into DNA LoBind tubes (Eppendorf 0030108.418).

#### Mus musculus RNA

Mouse embryonic stem (mES) E14 cells were maintained on plates coated with 0.2% gelatin in 2i/LIF medium at 37°C humidified incubator with 5% CO_2_. The culture medium consisted of a 1:1 mixture of DMEM/F12 (Thermo Fisher, #31331-028) and Neurobasal (Thermo Fisher, #12348-017), supplemented with 1% (v/v) penicillin–streptomycin (Sigma, #P4333), 2 μM GlutaMAX (Thermo Fisher, #35050-038), 0.1 mM non-essential amino acids (Thermo Fisher, #11140035), 1 mM sodium pyruvate (Thermo Fisher, #11360-039), 0.5x N-2 Supplement (Thermo Fisher, #17502-048), 0.5x B27 Supplement (Thermo Fisher, #17504-044), and 50 μM 2-mercaptoethanol (Thermo Fisher, #31350-038). The medium was further supplemented with 3 μM CHIR99021 (Sigma, #SML1046, GSK3 inhibitor), 1 μM PD0325901 (Sigma, #PZ0162, MEK inhibitor), and leukemia inhibitory factor (LIF-produced in-house). RNA was extracted using the TRIzol Plus RNA Purification Kit (Thermo 12183555) according to the manufacturer’s protocol and eluted into DNA LoBind tubes (Eppendorf 0030108.418).

#### Arabidopsis thaliana RNA

Whole 12-day-old seedlings of Columbia-0 wild-type *Arabidopsis thaliana* were harvested and ground into fine powder in liquid nitrogen. ~100 mg aliquot of the frozen tissue powder was made from each sample. RNA was extracted by TRI reagent and chloroform phase separation, precipitated with isopropanol, and washed with ice-cold 70% ethanol. In order to remove the excessive amount of polysaccharides from plant tissue, the pellet was resuspended in 10% ethanol and 0.1M NaOAc pH5.2, and RNA was precipitated again in 70% ethanol and 0.3M NaOAc pH5.2, washed with ice-cold 70% ethanol, air-dried and resuspended in nuclease-free water.

All RNA aliquots were stored at −80°C until use.

### Poly(A)+ RNA selection

Poly(A)+ RNA was enriched from both IVT spike-ins and transcriptome RNA samples using Dynabeads Oligo(dT)25 beads. RNA samples were adjusted with nuclease-free water, heated at 65°C for 2 min to disrupt secondary structure, and immediately placed on ice. Dynabeads were equilibrated in Binding Buffer before addition of RNA, and samples were mixed gently and incubated for 3–5 min at room temperature to allow hybridization to the oligo(dT) beads. Bead-bound RNA was washed twice with Washing Buffer B, with beads collected on a magnetic rack between washes. Poly(A)+ RNA was eluted in 10 mM Tris-HCl, pH 7.5 by heating at 70°C for 2 min, and the eluate was transferred to a fresh tube. Enriched RNA was quantified using the Qubit RNA High Sensitivity assay and assessed using an Agilent RNA Pico Bioanalyzer.

### Nanopore DRS library preparation and sequencing

Libraries were barcoded using either WarpDemuX (van der Toorn et al., 2025), then samples were pooled and prepared according to the SQK-RNA004 kit and sequenced for 72 hrs on a FLO-MIN004RA MinION flow cell, using the software MinKNOW. Similarly, separate libraries were barcoded SeqTagger barcodes (Pryszcz et al., 2024), and then samples were pooled and prepared according to the SQK-RNA004 kit and sequenced on a FLO-MIN004RA MinION flow cell.

### Data processing and creation of master table

Four direct RNA sequencing runs were performed (RNA004 chemistry; library preparation and sequencing described below), with the species, IVT and barcode assignments to each flowcell outlined in **Figure 1B**. After basecalling and read splitting the four runs yielded 17,747,356 reads in total (FC1: 1,813,929; FC2: 5,166,099; FC3: 5,170,214; FC4: 5,597,114).

Reads were basecalled using Dorado, and a master table was created containing read identifiers, flow-cell identifiers, mean RNA read quality, read length, aligned query length and poly(A)-tail length estimates. Read identities were obtained by aligning basecalled reads to a combined reference representing the four species (genomes hg38, GRCm39, TAIR10 and R64-1-1) and four IVTs (**Supplementary Table 1**), and appended to the master table. Demultiplexing outcomes were also appended to the table as described below.

### Demultiplexing calls

Three models were run: WarpDemuX WDX4 and WDX10 on FC1 and FC4, and SeqTagger b96_RNA004 on FC2 and FC3. Default comparisons used the final WarpDemuX calls at the nominal 99% settings and SeqTagger calls with barcode baseQ >=50. For threshold analyses, WarpDemuX confidence was defined as the difference between the highest and second-highest class scores. WDX4 analyses used the calls retained by its default workflow. WDX10 analyses also used provisional barcode predictions reconstructed from the exported class probabilities. Software versions and model identifiers are listed below, and commands and threshold tables are provided with the analysis scripts.

### Determination of ground truth read sets

Prior to barcode filtering, a read was included in the ground-truth set if it had a species or IVT reference identity with mapping quality (MAPQ) of at least 60 and at least 80% of its read length aligned to the reference. This yielded 12,317,897 species and 307,023 IVT reference reads, which remained fixed for all subsequent comparisons of precision and retention.

### Performance metrics for benchmarking

Reads were classified as in-panel if assigned to a loaded barcode, off-panel if assigned to an unused barcode included in the model, or not assigned if uncalled or not passing the method’s score filter. A correct assignment was defined as a reference read whose predicted barcode matched the barcode expected for that read’s origin. In-panel retention for a given origin was defined as the number of its reference reads assigned to any loaded barcode divided by all reference reads from that origin. In-panel precision for a barcode was defined as the number of correct assignments divided by all assignments to that barcode. Metrics were calculated separately for species and IVTs for each flow cell, unless otherwise described.

### Barcode-specific threshold comparisons

For precision–retention curves, all species reference reads were retained and weighted inversely to the number of reference reads from their species in that flow cell, giving each of the four species one-quarter of the total contribution. Precision was calculated from the weighted numbers of correct and total assignments to each barcode. Each curve varied the cutoff for one barcode while keeping assignments to the other barcodes at their default settings. We also calculated the same curves using the observed, unweighted read counts.

### Analysis of off-panel distributions

SeqTagger assignments to unused barcodes were examined at baseQ cutoffs of 40, 50 and 60. For each unused barcode, we calculated its assignment rate from each species, averaged these rates equally across species within a run, and then averaged the two runs equally. The resulting values were expressed as shares of all off-panel assignments and ranked to show which unused barcodes received the most calls. Observed read counts were also reported separately by run for SeqTagger and WDX10.

### Read quality analyses

Mean RNA read-quality scores and read lengths were extracted from the basecalled BAM files, and their distributions were plotted by flow cell. Reference reads were grouped into mean RNA quality bins of <10, 10–<15, 15–<20, 20–<25 and ≥25. In-panel retention and precision were calculated from all reference reads in each bin, separately for species and IVTs in each flow cell. These calculations used the observed unweighted read counts and points were shown when the relevant denominator contained at least 100 reads. Poly(A)-tail estimates were also examined to distinguish reads with no estimate from those with a positive estimated tail length.

### IVT based calibration and transfer to species

Cutoffs were selected using reads with IVT reference labels independently for each method and flow cell. In line with the published workflows, SeqTagger used one common cutoff across the four barcodes, whereas WarpDemuX used a specific cutoff for each barcode. For each precision target (99%, 99.5% or 99.9%), we selected the cutoff retaining the most IVT assignments while meeting the target and retaining at least 100 calls in each required group. The selected cutoffs were applied unchanged to species reads from the same flow cell (cis) or the other run of the same workflow (trans), matching cutoffs by barcode identity. Species reads were used to evaluate the selected settings, not to select them.

To assess the effect of IVT read count on calibration, 100 independent samples were drawn without replacement for each method and flow cell at sizes of 1,000, 2,000, 5,000, 10,000, 20,000 and 40,000 IVT reference reads. Off-panel, uncalled and unassigned reads were included in the sampling population.

For each sample, cutoffs were refitted at the 99% target using the same selection procedure. Cutoffs were evaluated on the complete species reference population from the same flow cell. The resulting distributions of precision and retention were used to assess calibration stability as the number of IVT reads increased.

### Final assignment yields

Default and 99% cis and trans IVT-calibrated cutoffs were applied to the complete read population, including reads outside the ground-truth reference sets. Total assignments to loaded barcodes were counted for each method. The R package eulerr was used to produce two-set Venn diagrams with proportional areas, based on pooling assigned reads for WDX4 and WDX10 across the two flow cells for each setting. The overlap counted reads assigned by both models, irrespective of whether their barcode calls matched.

## Software and tools

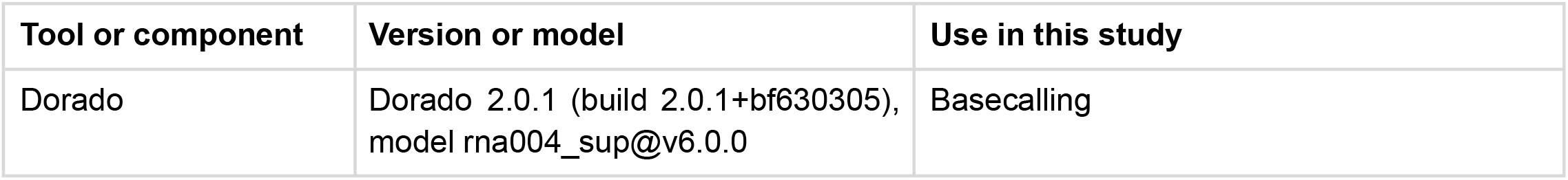
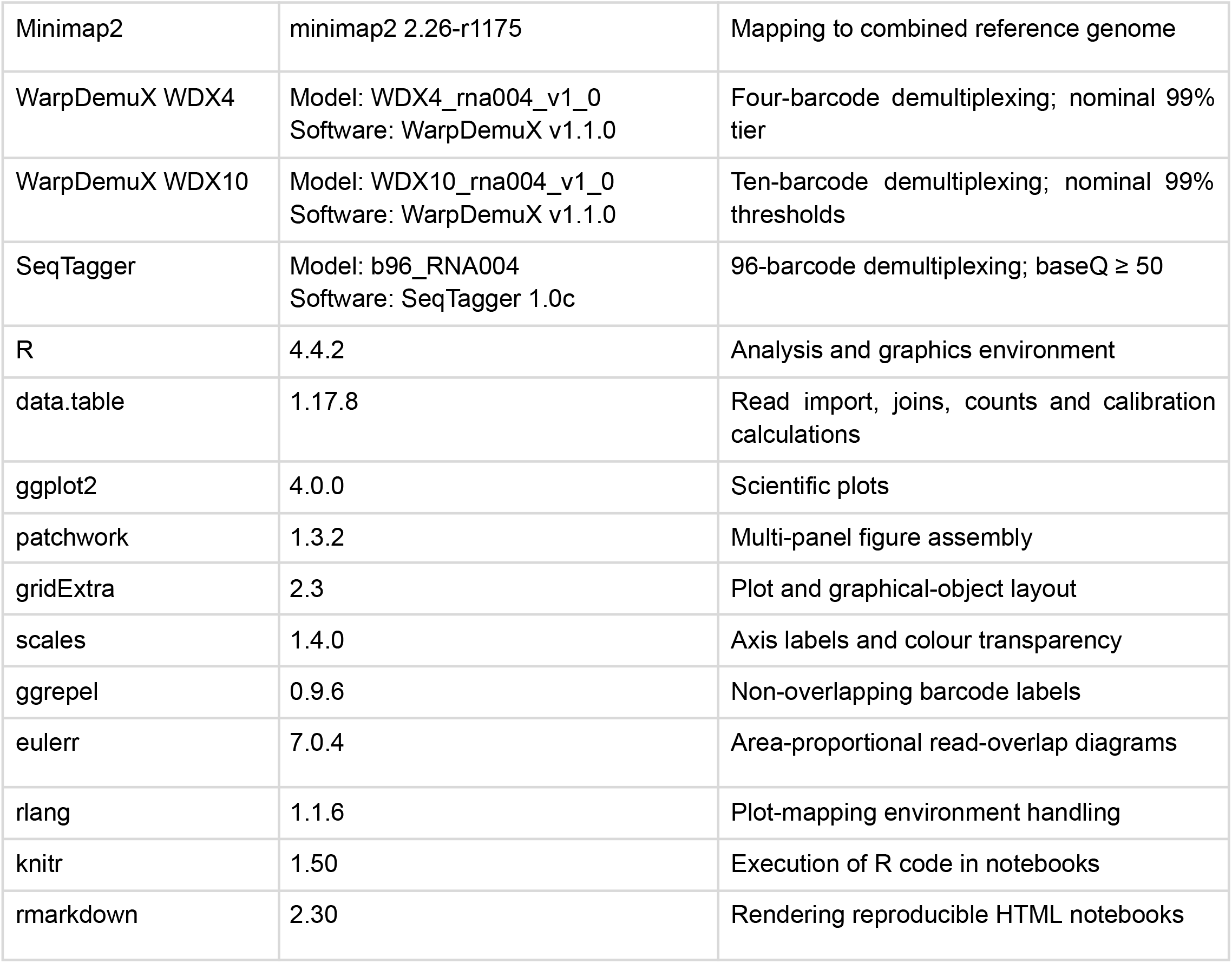

| Tool or component | Version or model | Use in this study |
| --- | --- | --- |
| Dorado | Dorado 2.0.1 (build 2.0.1+bf630305),<br>model rna004_sup@v6.0.0 | Basecalling |
| Minimap2 | minimap2 2.26-r1175 | Mapping to combined reference genome |
| WarpDemuX WDX4 | Model: WDX4_rna004_v1_0<br>Software: WarpDemuX v1.1.0 | Four-barcode demultiplexing; nominal 99% tier |
| WarpDemuX WDX10 | Model: WDX10_rna004_v1_0<br>Software: WarpDemuX v1.1.0 | Ten-barcode demultiplexing; nominal 99% thresholds |
| SeqTagger | Model: b96_RNA004<br>Software: SeqTagger 1.0c | 96-barcode demultiplexing; baseQ ≥ 50 |
| R | 4.4.2 | Analysis and graphics environment |
| data.table | 1.17.8 | Read import, joins, counts and calibration calculations |
| ggplot2 | 4.0.0 | Scientific plots |
| patchwork | 1.3.2 | Multi-panel figure assembly |
| gridExtra | 2.3 | Plot and graphical-object layout |
| scales | 1.4.0 | Axis labels and colour transparency |
| ggrepel | 0.9.6 | Non-overlapping barcode labels |
| eulerr | 7.0.4 | Area-proportional read-overlap diagrams |
| rlang | 1.1.6 | Plot-mapping environment handling |
| knitr | 1.50 | Execution of R code in notebooks |
| rmarkdown | 2.30 | Rendering reproducible HTML notebooks |

## Data Availability

Nanopore direct RNA sequencing data, including signal-level files, will be deposited in the European Nucleotide Archive (ENA), accession PRJEB127391. Code for reproducing the analyses will be available on GitHub (https://github.com/sarah-ku/drs_demultiplex_benchmark). The code relies on the master table containing annotations for all reads deposited on Zenodo, DOI: **10.5281/zenodo.22979500** (available upon final acceptance).

## Author contributions

S.R. and J.Y.A. conceived the project. J.B. and S.R. analysed the data. L.B.J. carried out the sequencing with support from J.B. J.Y.A. supervised the sequencing and carried out the yeast experiments, J.C. collected plant tissues and A.G. cultured the human and mouse cell lines. S.R. supervised the computational analyses. S.R. and J.B wrote the manuscript with input from all authors.

## Acknowledgements

We thank Professor Peter Brodersen, University of Copenhagen, Denmark for providing the seeds and facilities for *A. thaliana* experiments. We thank Dr. Amelie Stein, University of Copenhagen, Denmark, for providing *Saccharomyces cerevisiae* (BY4741).

## Supplementary figures

**Supplementary Figure 1:**
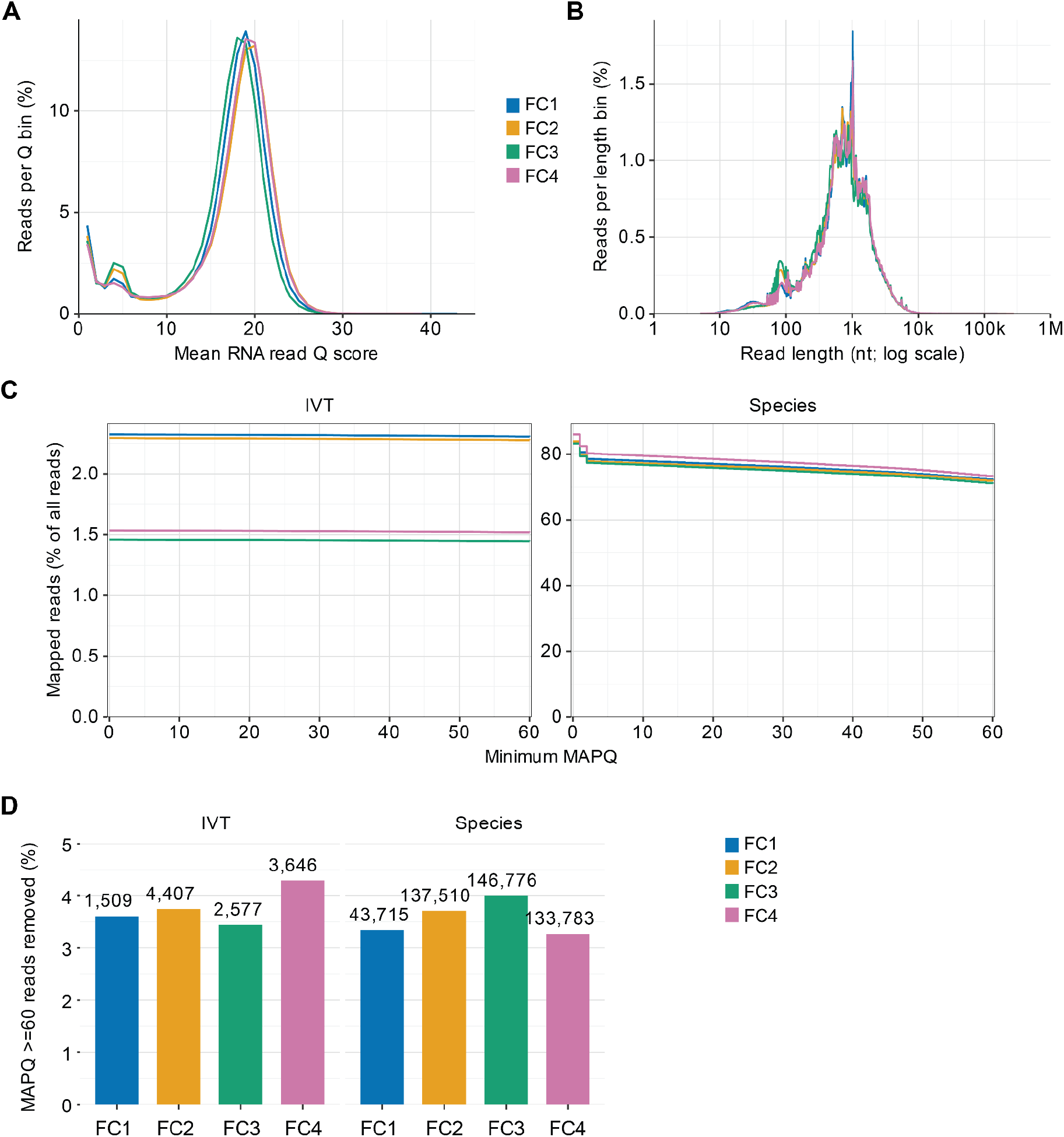
Quality control and mapping statistics. **(A)** Percentage of reads per mean RNA read quality bins, plotted for each flow cell. **(B)** Read-length distribution by flow cell. **(C)** Percentage of all reads mapping to IVT or species references, per MAPQ score threshold. IVTs on the left and species on the right. **(D)** Percentage of MAPQ60+ reads excluded due to an aligned read fraction below 80%.

**Supplementary Figure 2:**
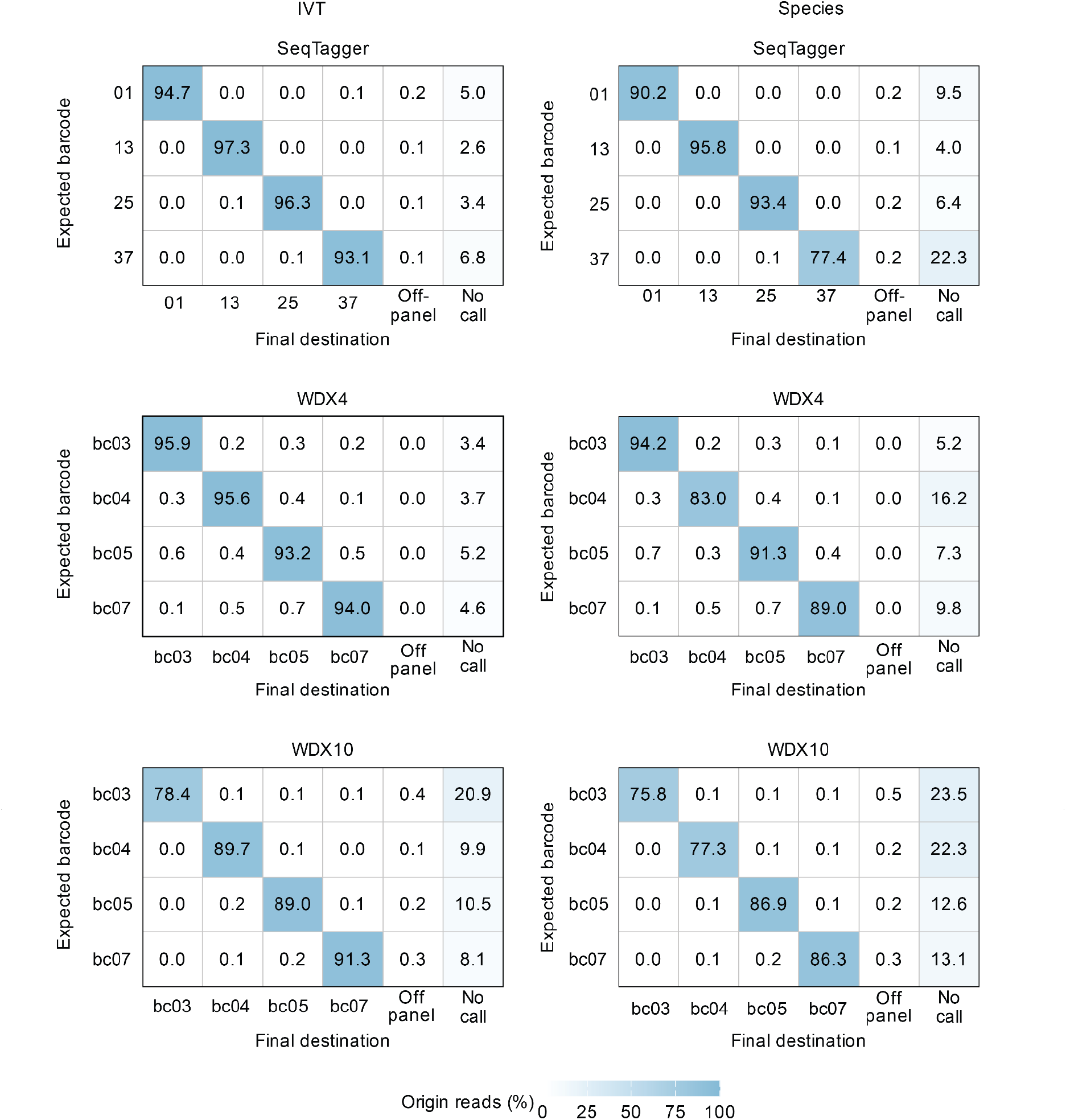
Confusion matrix showing barcode call allocation. For each of the three models, for both IVTs mapping reads only and species mapping reads, we show their barcode call rates for each of the IVTs/species, off-panel barcode calls (barcodes not used in the set) and reads where no call attempts were made. Calls are shown according to the recommended thresholds (Seqtagger: baseQ ≥50 filter, WarpDeMux: per barcode confidence scores for 99% precision target).

**Supplementary Figure 3:**
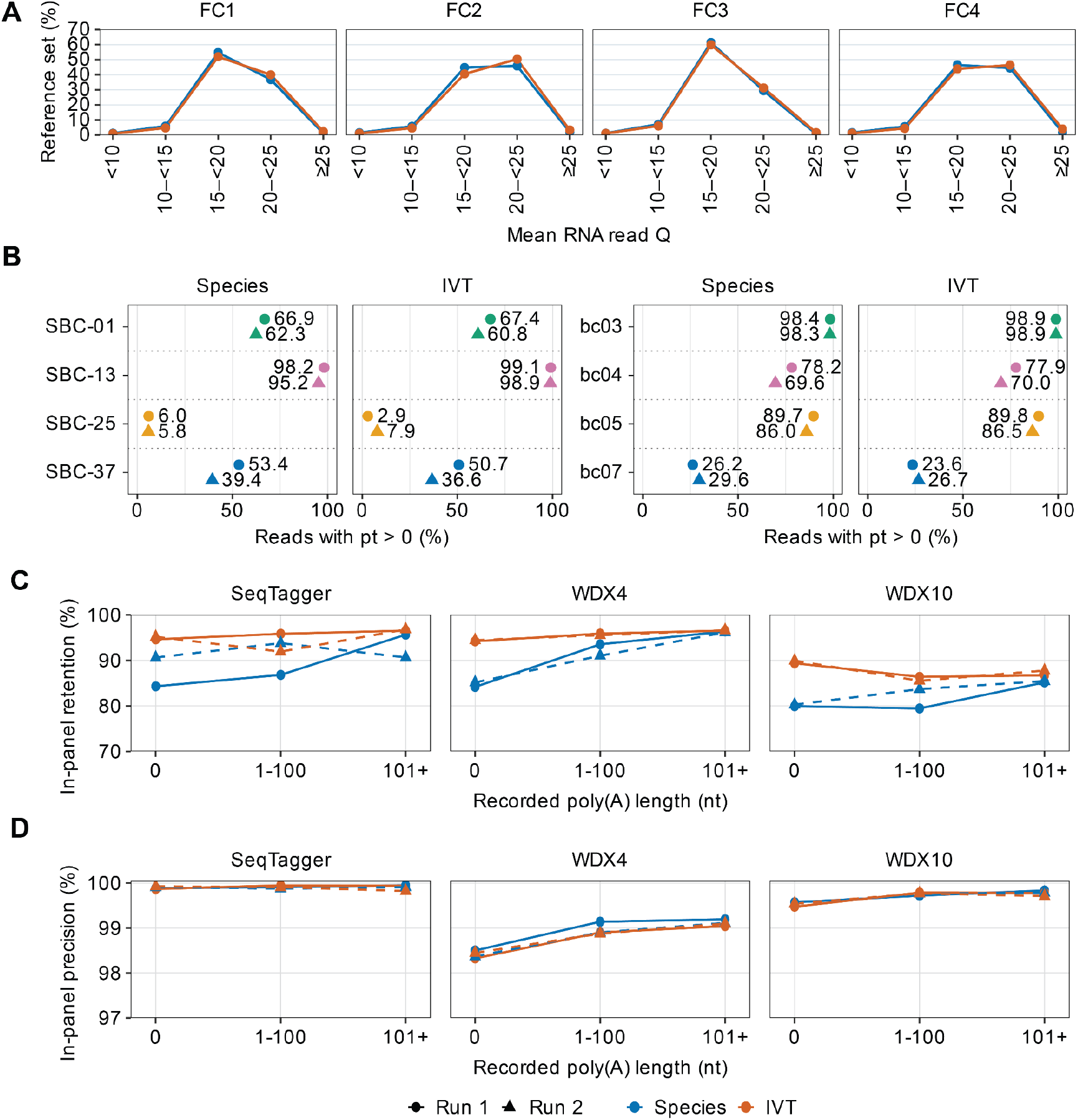
Investigation of causes of poor demultiplexing performance. **(A)** Percentage of reads in each reference set (IVT: orange, species: blue) falling into mean read quality score bins, split by flow cell. **(B)** Dorado calling rates for poly(A) tail, plotted as the percentage of reads with a non-zero tail length, plotted per run, method and for species and IVT. **(C, D)** In-panel retention and precision based on reads pooled according to method, run and type, split according to binned categories of Dorado-defined poly(A) tail length.

**Supplementary Figure 4:**
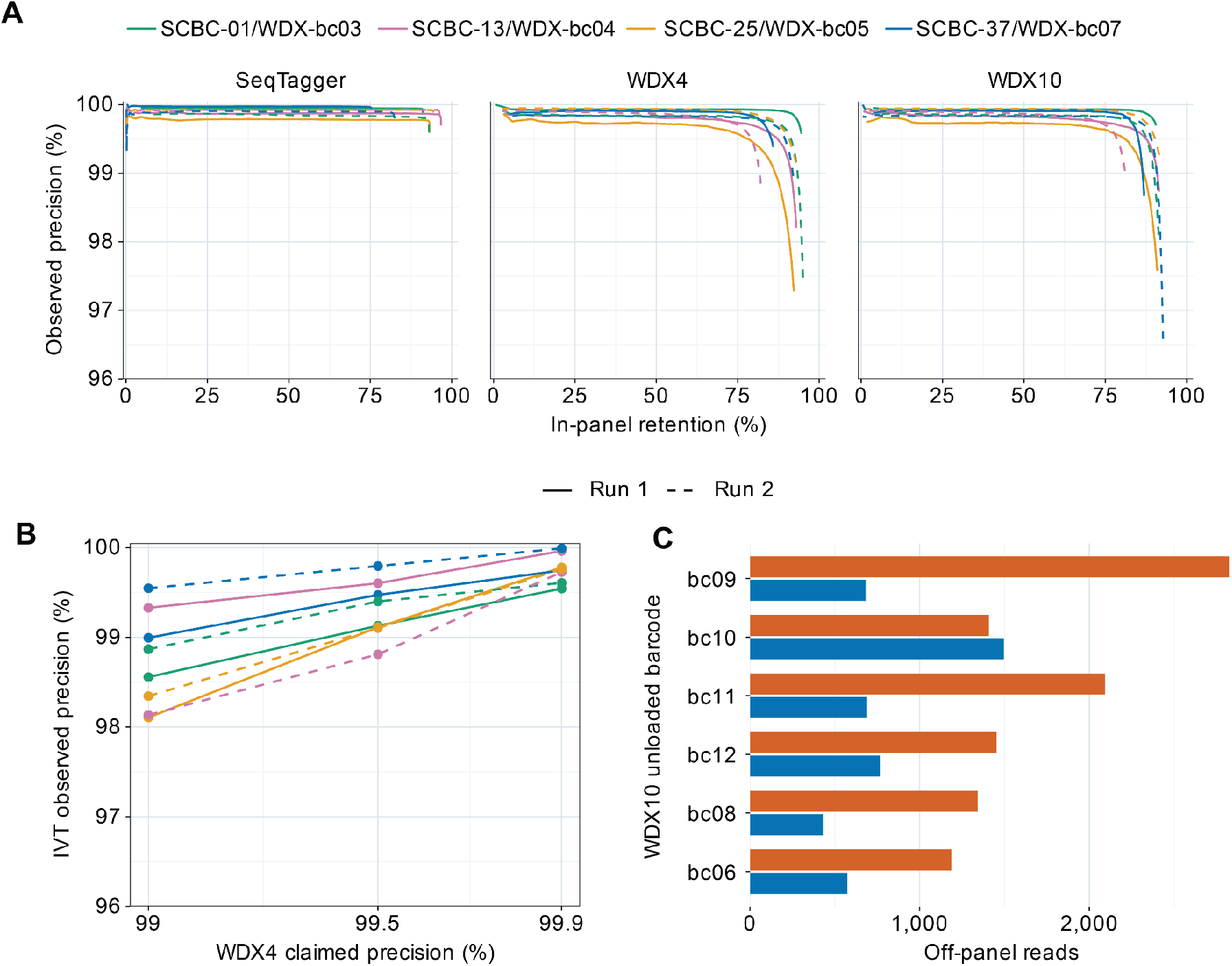
Precision and retention statistics according to varying thresholds. **(A)** as in Figure 4A but using unweighted counts. **(B)** Observed IVT precision at estimated shipped WDX4 tiers. Paired bars retain separate runs. **(C)** WDX10 off-panel counts at the native nominal-99% setting.

**Supplementary Figure 5:**
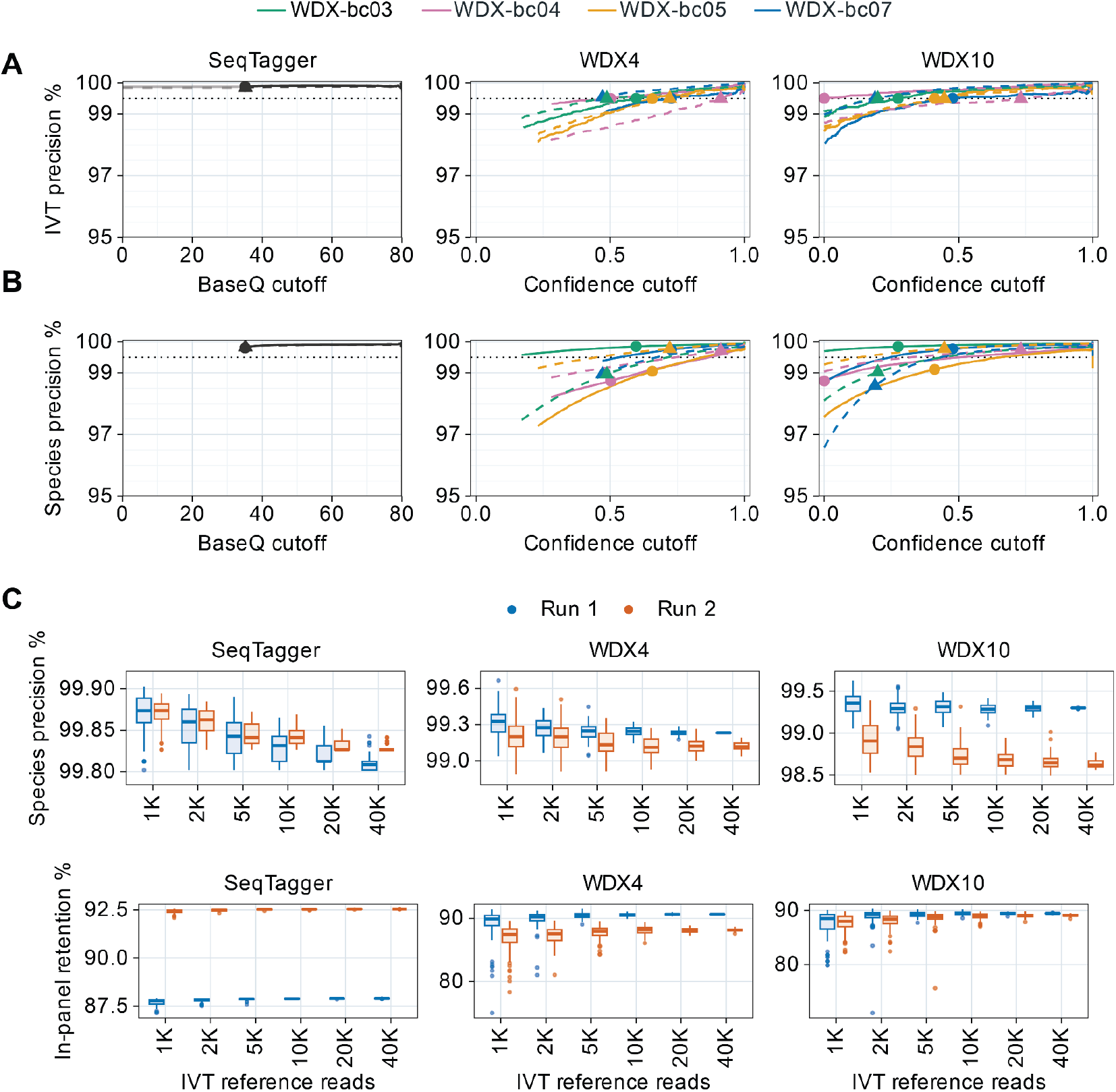
IVT based calibration. **(A,B)** as in Figure 5B,C, here optimised for the 99.5% precision target. **(C)** Number of IVT reads required for effective cis-based calibration. Points are based on resampling IVT read sets, calibrating and calculating precision and retention based on the species data within the same flow cell.

