## Supplementary Table 1 for "Systematic benchmark of demultiplexing workflows for direct RNA sequencing"

**Supplementary Table 1:** Sequences used for RNA spike-ins

**Renilla luciferase**

gctagccaccatgacttcgaaagtttatgatccagaacaaaggaaacggatgataactggtccgcagtggtgggccagatgtaaacaaatgaatg
ttcttgattcatttattaattattatgattcagaaaaacatgcagaaaatgctgttatttttttacatggtaacgcggcctcttcttatttatggcgacatgttgtgc
cacatattgagccagtagcgcggtgtattataccagaccttattggtatgggcaaatcaggcaaatctggtaatggttcttataggttacttgatcattac
aaatatcttactgcatggtttgaacttcttaatttaccaaagaagatcatttttgtcggccatgattggggtgcttgtttggcatttcattatagctatgagcatc
aagataagatcaaagcaatagttcacgctgaaagtgtagtagatgtgattgaatcatgggatgaatggcctgatattgaagaagatattgcgttgatc
aaatctgaagaaggagaaaaaatggttttggagaataacttcttcgtggaaaccatgttgccatcaaaaatcatgagaaagttagaaccagaaga
atttgcagcatatcttgaaccattcaaagagaaaggtgaagttcgtcgtccaacattatcatggcctcgtgaaatcccgttagtaaaaggtggtaaacc
tgacgttgtacaaattgttaggaattataatgcttatctacgtgcaagtgatgatttaccaaaaatgtttattgaatcggacccaggattcttttccaatgcta
ttgttgaaggtgccaagaagtttcctaatactgaatttgtcaaagtaaaaggtcttcatttttcgcaagaagatgcacctgatgaaatgggaaaatatatc
aaatcgttcgttgagcgagttctcaaaaatgaacaataattctagactcgagccggaagttttc

**Firefly luciferase**

GAGACCCAAGCTTATGCATGCGGCCGCATCTAGAGGGCCCGGATCCAAATGGAAGACGCCAAAAAC
ATAAAGAAAGGCCCGGCGCCATTCTATCCTCTAGAGGATGGAACCGCTGGAGAGCAACTGCATAAG
GCTATGAAGAGATACGCCCTGGTTCCTGGAACAATTGCTTTTACAGATGCACATATCGAGGTGAACA
TCACGTACGCGGAATACTTCGAAATGTCCGTTCGGTTGGCAGAAGCTATGAAACGATATGGGCTGAA
TACAAATCACAGAATCGTCGTATGCAGTGAAAACTCTCTTCAATTCTTTATGCCGGTGTTGGGCGCGT
TATTTATCGGAGTTGCAGTTGCGCCCGCGAACGACATTTATAATGAACGTGAATTGCTCAACAGTATG
AACATTTCGCAGCCTACCGTAGTGTTTGTTTCCAAAAAGGGGTTGCAAAAAATTTTGAACGTGCAAAA
AAAATTACCAATAATCCAGAAAATTATTATCATGGATTCTAAAACGGATTACCAGGGATTTCAGTCGAT
GTACACGTTCGTCACATCTCATCTACCTCCCGGTTTTAATGAATACGATTTTGTACCAGAGTCCTTTG
ATCGTGACAAAACAATTGCACTGATAATGAATTCCTCTGGATCTACTGGGTTACCTAAGGGTGTGGC
CCTTCCGCATAGAACTGCCTGCGTCAGATTCTCGCATGCCAGAGATCCTATTTTTGGCAATCAAATCA
TTCCGGATACTGCGATTTTAAGTGTTGTTCCATTCCATCACGGTTTTGGAATGTTTACTACACTCGGA
TATTTGATATGTGGATTTCGAGTCGTCTTAATGTATAGATTTGAAGAAGAGCTGTTTTTACGATCCCTT
CAGGATTACAAAATTCAAAGTGCGTTGCTAGTACCAACCCTATTTTCATTCTTCGCCAAAAGCACTCT
GATTGACAAATACGATTTATCTAA

**mRFP1**

ATGGCTTCCTCCGAAGACGTTATCAAAGAGTTCATGCGTTTCAAAGTTCGTATGGAAGGTTCCGTTAA
CGGTCACGAGTTCGAAATCGAAGGTGAAGGTGAAGGTCGTCCGTACGAAGGTACCCAGACCGCTAA
ACTGAAAGTTACCAAAGGTGGTCCGCTGCCGTTCGCTTGGGACATCCTGTCCCCGCAGTTCCAGTA
CGGTTCCAAAGCTTACGTTAAACACCCGGCTGACATCCCGGACTACCTGAAACTGTCCTTCCCGGAA
GGTTTCAAATGGGAACGTGTTATGAACTTCGAAGACGGTGGTGTTGTTACCGTTACCCAGGACTCCT
CCCTGCAAGACGGTGAGTTCATCTACAAAGTTAAACTGCGTGGTACCAACTTCCCGTCCGACGGTCC
GGTTATGCAGAAAAAAACCATGGGTTGGGAAGCTTCCACCGAACGTATGTACCCGGAAGACGGTGC
TCTGAAAGGTGAAATCAAAATGCGTCTGAAACTGAAAGACGGTGGTCACTACGACGCTGAAGTTAAA
ACCACCTACATGGCTAAAAAACCGGTTCAGCTGCCGGGTGCTTACAAAACCGACATCAAACTGGACA
TCACCTCCCACAACGAAGACTACACCATCGTTGAACAGTACGAACGTGCTGAAGGTCGTCACTCCAC
CGGTGCTTAA

**AmpR (ampicillin resistance gene)**

atgagtattcaacatttccgtgtcgcccttattcccttttttgcggcattttgccttcctgtttttgctcacccagaaacgctggtgaaagtaaaagatgctgaa
gatcagttgggtgcacgagtgggttacatcgaactggatctcaacagcggtaagatccttgagagttttcgccccgaagaacgttttccaatgatgag
cacttttaaagttctgctatgtggcgcggtattatcccgtattgacgccgggcaagagcaactcggtcgccgcatacactattctcagaatgacttggttg
agtactcaccagtcacagaaaagcatcttacggatggcatgacagtaagagaattatgcagtgctgccataaccatgagtgataacactgcggcc
aacttacttctgacaacgatcggaggaccgaaggagctaaccgcttttttgcacaacatgggggatcatgtaactcgccttgatcgttgggaaccgg
agctgaatgaagccataccaaacgacgagcgtgacaccacgatgcctgtagcaatggcaacaacgttgcgcaaactattaactggcgaactactt
actctagcttcccggcaacaattaatagactggatggaggcggataaagttgcaggaccacttctgcgctcggcccttccggctggctggtttattgct
gataaatctggagccggtgagcgtgggtctcgcggtatcattgcagcactggggccagatggtaagccctcccgtatcgtagttatctacacgacgg
ggagtcaggcaactatggatgaacgaaatagacagatcgctgagataggtgcctcactgattaagcattggtaa

**Supplementary Table 2:** Demultiplexing cutoffs used in Figure 5

Default settings and cutoffs selected from IVT reference reads at minimum empirical precision targets of 99%, 99.5% and 99.9%.

| Method | IVT calibration run / flow cell | Barcode | Default | IVT target 99% | IVT target 99.5% | IVT target 99.9% |
|---|---|---|---|---|---|---|
| SeqTagger | 1 / FC2 | All loaded | 50 | 35.1 | 35.1 | 37.3 |
| SeqTagger | 2 / FC3 | All loaded | 50 | 35.2 | 35.2 | 75.1 |
| WDX4 | 1 / FC1 | BC03 | 0.17 | 0.341 | 0.596 | 0.994 |
| WDX4 | 1 / FC1 | BC04 | 0.28 | 0.281 | 0.501 | 0.941 |
| WDX4 | 1 / FC1 | BC05 | 0.23 | 0.478 | 0.656 | 0.998 |
| WDX4 | 1 / FC1 | BC07 | 0.47 | 0.476 | 0.727 | 1.000 |
| WDX4 | 2 / FC4 | BC03 | 0.17 | 0.220 | 0.487 | 0.990 |
| WDX4 | 2 / FC4 | BC04 | 0.28 | 0.702 | 0.911 | 0.999 |
| WDX4 | 2 / FC4 | BC05 | 0.23 | 0.466 | 0.721 | 0.980 |
| WDX4 | 2 / FC4 | BC07 | 0.47 | 0.471 | 0.471 | 0.847 |
| WDX10 | 1 / FC1 | BC03 | 0.86 | 0.0208 | 0.2751 | 0.9651 |
| WDX10 | 1 / FC1 | BC04 | 0.59 | 0.0009 | 0.0009 | 0.7537 |
| WDX10 | 1 / FC1 | BC05 | 0.40 | 0.1723 | 0.4112 | 0.9967 |
| WDX10 | 1 / FC1 | BC07 | 0.49 | 0.1852 | 0.4787 | 0.9904 |
| WDX10 | 2 / FC4 | BC03 | 0.86 | 0.0006 | 0.2000 | 0.9585 |
| WDX10 | 2 / FC4 | BC04 | 0.59 | 0.1412 | 0.7322 | 0.9924 |
| WDX10 | 2 / FC4 | BC05 | 0.40 | 0.1492 | 0.4476 | 0.9749 |
| WDX10 | 2 / FC4 | BC07 | 0.49 | 0.0046 | 0.1893 | 0.6696 |
